# *In silico* characterization, homology modeling and structure-based functional annotation of Nile Tilapia (*Oreochromis niloticus*) Hsp70 and Hsc70 proteins

**DOI:** 10.1101/2023.12.27.573401

**Authors:** Geraldine B. Dayrit, Normela Patricia F. Burigsay, Emmanuel M. Vera Cruz, Mudjekeewis D. Santos

## Abstract

**Background:** The molecular chaperones known as heat shock proteins 70 (Hsp70) and heat shock cognate protein 70 (Hsc70) are vital for maintaining cellular integrity and controlling stress.

**Methodology:** The *On-*Hsp70 and *On-*Hsc70 proteins from Nile tilapia (*Oreochromis niloticus*) have been thoroughly examined in this study using *in silico* analysis, homology modeling, and functional annotation. Homology modeling was carried out using the SWISS-MODEL program, and the proposed model was assessed for its high reliability through analyses including ProSA, Verify 3D, PROVE, ERRAT, and Ramachandran plot.

**Results:** The essential features of the *On-*Hsp70 and *On-*Hsc70 proteins encompass amino acid lengths (640 and 645), molecular weights (70,233.48 and 70,773.17 Da), theoretical isoelectric points (pI = 5.63 and 5.28), and the overall counts of negatively and positively charged residues (95 and 86; 95 and 81). Furthermore, the instability index (II) values were 35.27 (*On-*Hsp70) and 38.85 (*On-*Hsc70). Similarly, the aliphatic index (AI) exhibited high values for both proteins, reaching 84.58 (*On-*Hsp70) and 82.85 (*On-*Hsc70). *On-*Hsp70 and *On-*Hsc70 were both shown to contain an MreB/Mbl domain.

**Discussion:** The authors found that *On-*Hsp70 and *On-*Hsc70 share key characteristics, including an acidic nature, high stability, and conserved domains. Protein-protein interaction analysis identified the co-chaperone Stip1 as a primary functional partner. Comparative modeling yielded highly reliable 3D models, revealing structural similarity to known proteins and predicted binding sites. Furthermore, the primary subcellular localization of both proteins is the cytoplasm. Functional analysis predicted an AMP-PNP binding site for *On-*Hsp70 and ATP binding site for *On-*Hsc70.

**Conclusion:** The discoveries deepen our understanding of Hsc70 and Hsp70 in Nile tilapia, highlighting their importance in fish physiology and positioning them as crucial study topics moving forward. This study adds to our understanding of the actions of these proteins in cellular processes and stress responses, which could impact fish health and resilience.

## Introduction

In addition to being a popular and important fish species for the economy, tilapia (*Oreochromis* spp.) is also a topic of great interest in molecular research, especially with regard to heat shock proteins (Hsp). In many different organisms, including fish, these molecular chaperones are essential for protein folding, cellular integrity, and stress responses (Hu et al., 2022). Extensive documentation of Hsp in fish underscores the significant role in addressing stress and preserving the integrity of cells (Roberts et al., 2010). A study by Zhang et al. (2014) discovered four Hsps from the Hsp70 family that exhibit conserved properties in their amino acid sequences in Nile tilapia (*O. niloticus*). Comprehending the complex processes involving Hsp genes is essential to understanding how tilapia respond to environmental stressors and preserve cellular homeostasis.

Hsp70 and its constitutively expressed counterpart, heat shock cognate protein 70 (Hsc70), are key members of the Hsp family. Hsp70 and Hsc70 function similarly and share 85% of their identity in humans (Liu et al., 2012). Further, B-cell lymphoma-2 (BCL2) associated Athano-Gene 1 (BAG-1) binds the ATPase domains of Hsp70 and Hsc70, modulating their chaperone activity and functioning as a competitive antagonist of the co-chaperone Hip.

Hsp70/Hsc70 chaperone capabilities are defined and diversified by interactions with different BAG-family proteins (Takayama et al., 1999). These proteins become even more interesting when considering fish. Fish are ectothermic animals, making them especially vulnerable to changes in temperature and other environmental stresses (Volkoff and Rønnestad, 2020). Heat shock protein 70 are well known for their conceivable function in temperature adaptation and are essential for cellular thermotolerance in fish (Wang et al., 2020). While the heat shock cognate 71-kDa protein (Hsc70) is the molecular counterpart that is constitutively expressed (Silva et al., 2021). In a variety of cellular activities, including protein synthesis, folding, transport across membrane channels, translocation, and denaturation, the Hsc70 protein functions as a molecular chaperone (Tran et al., 2015).

Under stress, a stress-inducible Hsp70 gene and the “housekeeping” Hsc 70 gene are the two distinct genes that are primarily linked to physiological functions and encode the Hsp70 family.

By examining both the structural and functional aspects of Hsp70 and Hsc70 in Nile tilapia, this study seeks to provide a comprehensive understanding of these proteins. Our specific goal is to determine the specific distinctions between Hsp70 and Hsc70 in terms of physicochemical characteristics, structure, subcellular localization, and protein interactions. By tackling this research question, we may be able to gain new insights into the adaptations of tilapia and illuminate the basic molecular mechanisms regulating their cellular functions and stress responses.

Studying Hsp70 and Hsc70 in Nile tilapia holds significant importance. These proteins are instrumental in the fish’s ability to withstand stressors, such as temperature fluctuations, heavy metals, oxidative stress, and infections, which are prevalent in aquaculture and natural environments (Jeyachandran, 2023). To the best of our knowledge, no in silico published three-dimensional (3D) models of proteins from the *O. niloticus* family have been made.

Understanding the structural and functional nuances of these proteins in tilapia can provide valuable insights for enhancing stress management strategies in aquaculture, optimizing fish welfare, and even contributing to the broader field of cellular biology. Furthermore, insights gained from this study may have applications beyond tilapia and extend to other fish species, offering potential benefits in fisheries and aquatic ecosystems.

In the forthcoming sections of this paper, we will present the methodologies employed for our in-depth analysis of tilapia’s Hsp70 and Hsc70 proteins. This study integrates physicochemical characterization, domain architecture analysis, comparative homology modeling, structural analysis, and functional annotation to provide a comprehensive overview of these proteins. The results will be discussed in the context of their implications for stress responses and cellular functions, not only in Nile Tilapia but also in the broader domain of fish physiology and potentially in the wider spectrum of molecular chaperone research.

## Materials & Methods

### Sequence retrieval

The mRNA sequences of *On-*Hsp70 (Accession No.: XM_019357557.1) and *On-*Hsc70 (Accession No.: WLP61716.1) were uploaded to Expasy’s Translate Server with the following format: compact formatting (Compact: M) with no spaces on both the forward and reverse strands (Galsteiger et al., 2005). Afterward, the obtained protein sequences were aligned with each other using Clustal Omega, a tool available on the EMBL-EBI server, in accordance with the methodology outlined by McWilliam et al. (2013). Percent identity was then calculated using BLAST (Atschul et al., 1997).

### Physicochemical characterization

The amino acid sequences of Hsp70 and Hsc70 from *O. niloticus* were submitted to Expasy’s ProtParam Server to determine their physicochemical properties (Galsteiger et al., 2005). A protein sequence or Swiss-Prot or TrEMBL entry can be used to compute physical and chemical parameters for the protein.

### Domain Architecture Analysis

The conserved domains found in the Hsp70 and Hsc70 were identified using domain architecture analysis. This was conducted utilizing SMART (Letunic et al., 2021) to study simple molecular architecture. Also, this allows the identification and annotation of genetically mobile domains and the analysis of domain architectures. These domains are thoroughly documented concerning their presence across different species, their functional categories, their three-dimensional structures, and the critical residues essential for their functions. Every domain discovered in a non*-*repetitive protein database, along with the search criteria and taxonomic details, was meticulously recorded in a relational database system (Schultz et al., 2000).

### Comparative homology modeling

The SWISS-MODEL service was used to model the proteins’ homology. This service generates a set of projected models by aligning the input target protein with existing templates (Arnold et al., 2006). Sequence identity and GMQE were used to choose the best template for building the 3D model (Fiser, 2004). GalaxyRefine was then used to refine protein structure. This method starts with side-chain reconstruction, then side-chain repacking, and finally relaxation of the entire structure via molecular dynamics simulations (Heo et al., 2013). Next, several analyses, including PROCHECK’s Ramachandran plot analysis, ERRAT, PROVE, Verify3D, and ProSA (all accessible from the SAVES server at http://nihserver.mbi.ucla.edu), were used to evaluate the stereochemical quality and correctness of the predicted models (Sippl et al., 1993). Additionally, structural analysis was carried out, and Swiss PDB Viewer was used to create model representations.

### Subcellular localization

To predict the proteins’ subcellular position, the DeepLoc-1.0 server was used (Almagro Armenteros et al., 2017). The Neural Networks algorithm used by this server was trained on eukaryotic proteins from UniProt that have been known to have experimental evidence supporting their subcellular localization. It was important to note that this prediction process bases all conclusions entirely on information from the protein’s sequence.

### Structural similarity and functional annotation

The TM-align method was used to perform a global structural match on the COFACTOR web server. The TM-score, a measure of the overall structural similarity, was calculated as a result of this comparison. A TM-score of 1.0 indicates a perfect match between two structures (Zhang and Skolnick, 2005). TM-scores range from 0.0 to 1.0. Scores below 0.17, on the other hand, indicate that the proteins were selected at random and are unrelated. A score higher than 0.5 denotes that the structures have a fold that is usually similar (Zhang, 2008). The I-TASSER suite was used to annotate the analysis with regard to ligand-binding locations, gene ontology, and enzyme commission (Yang et al., 2015).

## Results

### Sequence retrieval

Figure 1 displays the sequence alignment of *On-*Hsp70 and *On-*Hsc70. BLAST results showed an 84.23% identity between these two heat shock proteins.

**Figure 1.**
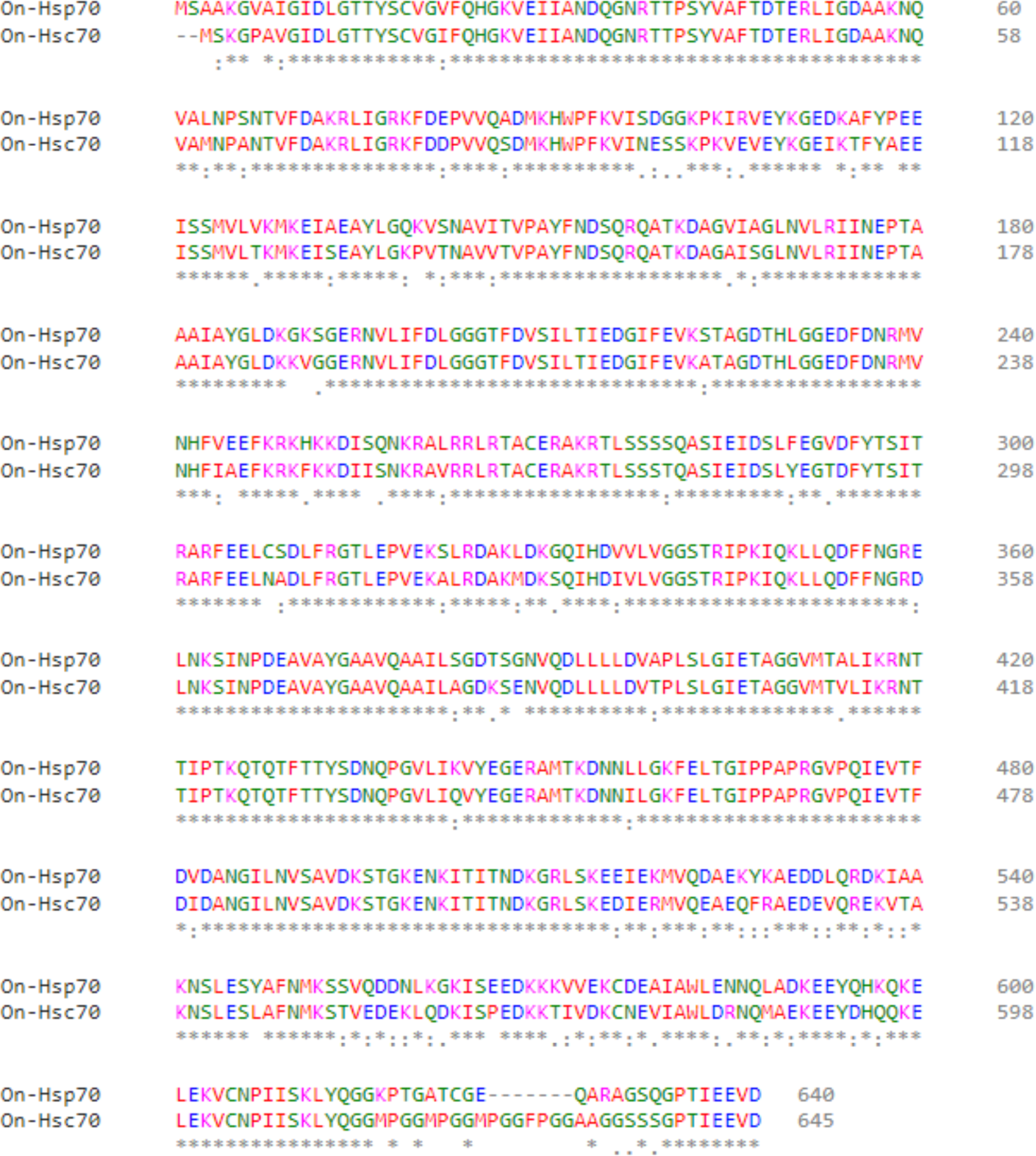
Alignment of *On*-Hsp70 and *On*-Hsc70 showed an 84.32% percent identity.

### Physicochemical Characteristics

The key characteristics of the *On-*Hsp70 and *On-*Hsc70 protein are summarized in Table 1 including amino acid lengths (640 and 645), molecular weights (70,233.48 and 70,773.17 Da), theoretical isoelectric points (pI = 5.63 and 5.28), and the total counts of negatively and positively charged residues (95 and 86; 95 and 81). Additionally, the calculated extinction coefficient (EC), directly linked to the cysteine, tryptophan, and tyrosine content, at 280 nm absorbance, amounted to 33725/32110 (assuming all pairs of cysteine residues form cysteines) and 33350/31860 (assuming all cysteine residues are reduced) M−1 cm−1 for *On-*Hsp70 and *On-*Hsc70 proteins, respectively. Furthermore, the instability index (II) values were 35.27 (*On-*Hsp70) and 38.85 (*On-*Hsc70); while the aliphatic index (AI) exhibited high values for both proteins, reaching 84.58 (*On-*Hsp70) and 82.85 (*On-*Hsc70).

**Table 1.** Physicochemical properties of *On-*Hsp70 and *On-*Hsc70.

| Property | On-Hsp70 | On-Hsc70 |
| --- | --- | --- |
| Number of amino acids | 640 | 645 |
| Theoretical pI | 5.63 | 5.28 |
| Amino Acid Composition (% of total amino acids) |  |  |
| Ala (A) | 53 (8.3%) | 50 (7.8%) |
| Arg (R) | 28 (4.4%) | 29 (4.5%) |
| Asn (N) | 31 (4.8%) | 32 (5.0%) |
| Asp (D) | 46 (7.2%) | 45 (7.0%) |
| Cys (C) | 6 (0.9%) | 4 (0.6%) |
| Gln (Q) | 27 (4.2%) | 25 (3.9%) |
| Glu (E) | 49 (7.7%) | 50 (7.8%) |
| Gly (G) | 51 (8.0%) | 53 (8.2%) |
| His (H) | 7 (1.1%) | 6 (0.9%) |
| Ile (I) | 44 (6.9%) | 48 (7.4%) |
| Leu (L) | 47 (7.3%) | 42 (6.5%) |
| Lys (K) | 58 (9.1%) | 52 (8.1%) |
| Met (M) | 9 (1.4%) | 15 (2.3%) |
| Phe (F) | 23 (3.6%) | 25 (3.9%) |
| Pro (P) | 21 (3.3%) | 26 (4.0%) |
| Ser (S) | 38 (5.9%) | 36 (5.6%) |
| Thr (T) | 39 (6.1%) | 45 (7.0%) |
| Trp (W) | 2 (0.3%) | 2 (0.3%) |
| Tyr (Y) | 15 (2.3%) | 14 (2.2%) |
| Val (V) | 46 (7.2%) | 46 (7.1%) |
| Pyl (O) | 0 (0.0%) | 0 (0.0%) |
| Sec (U) | 0 (0.0%) | 0 (0.0%) |
| (B) | 0 (0.0%) | 0 (0.0%) |
| (Z) | 0 (0.0%) | 0 (0.0%) |
| (X) | 0 (0.0%) | 0 (0.0%) |
| Total number of negatively charged residues (Asp + Glu) | 95 | 95 |
| Total number of positively charged residues (Arg + Lys) | 86 | 81 |
| Ext. coefficient (assuming all pairs of Cys residues form cystines) | 33,725 | 32,110 |
| Ext. coefficient (assuming all Cys residues are reduced) | 33,350 | 31,860 |

The estimated half-life values were consistent for both proteins: exceeding 30 hours in mammalian reticulocytes (in vitro), over 20 hours in yeast (in vivo), and more than 10 hours in Escherichia coli (in vivo). Furthermore, the grand average hydropathicity (GRAVY) of *On-*Hsp70 and *On-*Hsc70 was found to be -0.440 and -0.404, respectively. An analysis of amino acid composition revealed that *On-*Hsp70 had elevated levels of alanine (8.3%), lysine (9.1%), and glycine (8.0%), while *On-*Hsc70 contained higher proportions of alanine (7.8%), glutamic acid (7.8%), lysine (8.1%), and glycine (8.2%).

### Domain Architecture Analysis

Figure 2 shows *On-*Hsp70 and *On-*Hsc70 containing an MreB/Mbl domain at positions 119 to 383 and 108 to 382, respectively. The only difference detected between *On-*Hsp70 and *On-* Hsc70 was the low complexity region along positions 613 to 639 on *On-*Hsc70. The amino acid sequences known as low-complexity regions (LCRs), which are extremely common in eukaryotic proteins, contain repeats of one amino acid or brief amino acid motifs (Kastano et al., 2021).

**Figure 2.**
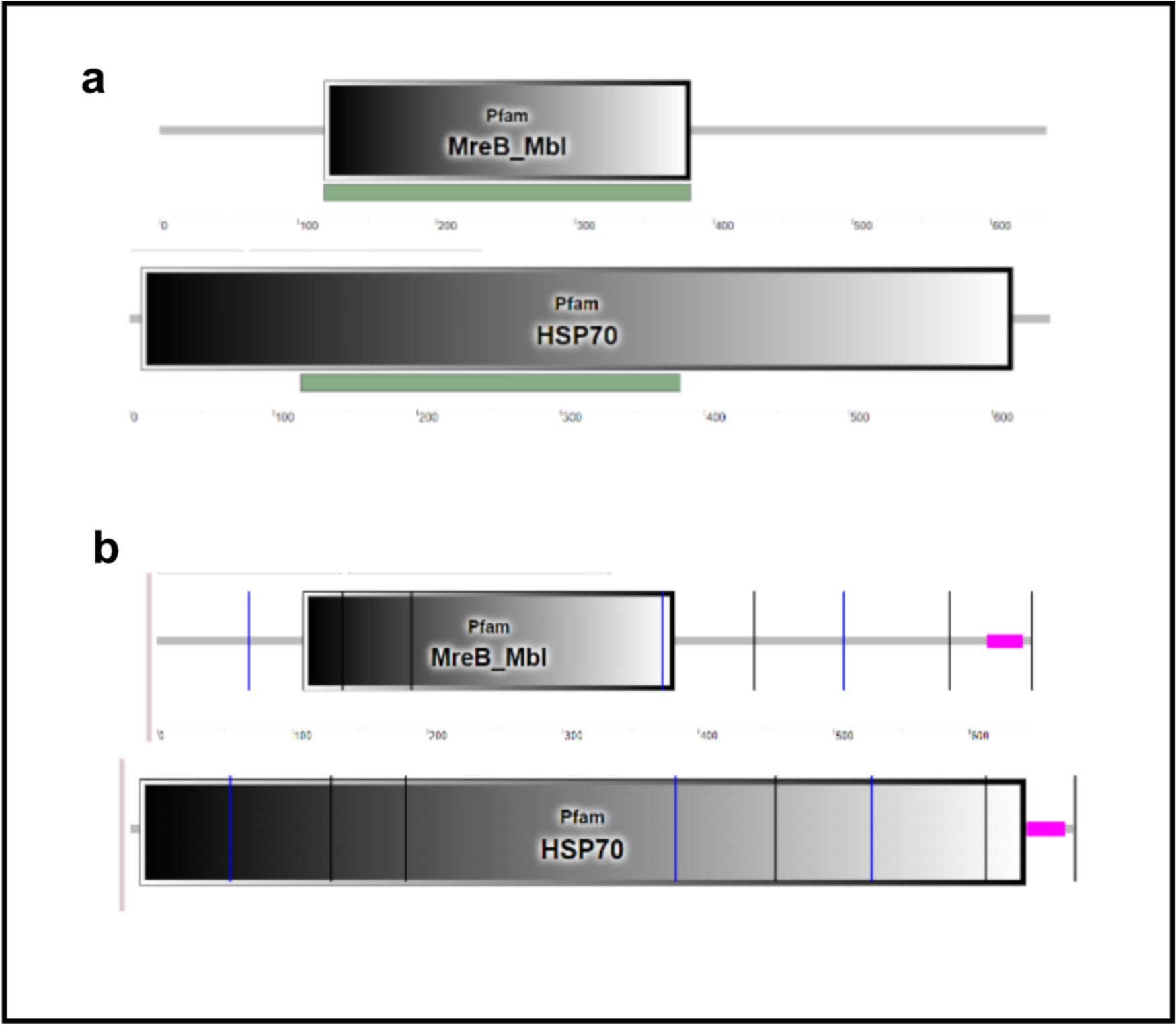
Domain architecture of *On*-Hsp70 and *On*-Hsc70.

### Comparative homology modeling

The hsp 70A Ciliary C1 central pair apparatus from *Chlamydomonas reinhardtii* (PDB ID: 7sqc) was selected as the template for homology modeling of *On-*Hsp70 and *On-*Hsc70 exhibited in Fig. 3, showing sequence identities of 74.10% and 74.65%, respectively.

**Figure 3.**
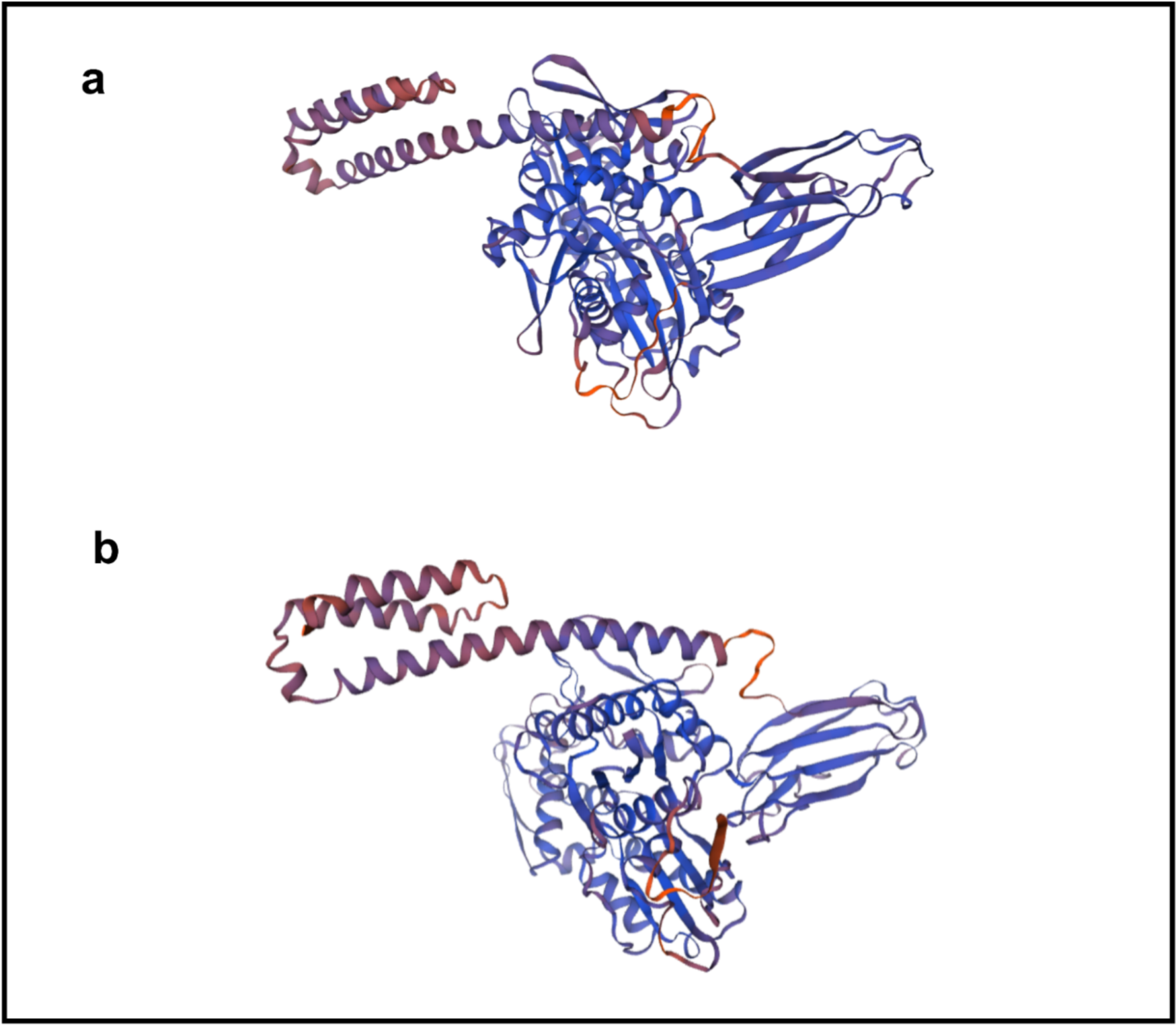
Predicted models of *On*-Hsp70 (a) and *On*-Hsc70 (b)

The quality of the models was assessed using various validation tools summarized in Table 2, while Fig. 4 reveals their corresponding plots. The Ramachandran plot analysis revealed excellent quality, with 95.50% of residues falling within the most favored regions for *On-*Hsp70 and 95.40% for *On-*Hsc70. The overall G-factor values for *On-*Hsp70 (0.02) and *On-*Hsc70 (0.20) were well within the acceptable range, indicating the reliability of the models.

**Figure 4.**
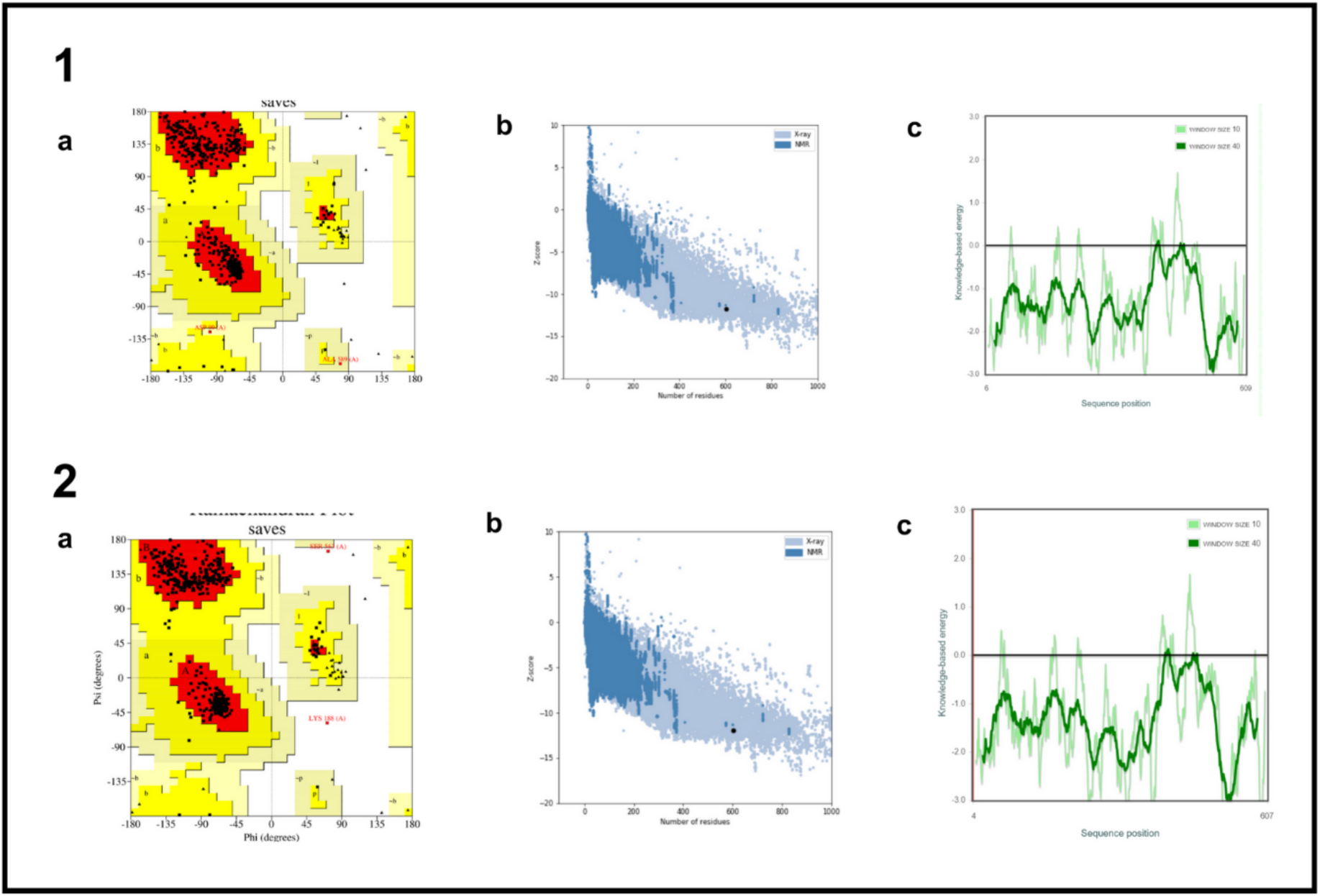
Validation indices for 1 *On*-Hsp70 and 2 *On*-Hsc70. a Ramachandran plot analysis shows where residues are located, emphasizing those that are in the desirable (red), permissible (yellow), liberally allowed (light yellow), and forbidden (white) categories. b Plotted alongside Z-scores from several experimentally determined protein chains in the Protein Data Bank is the model’s Z-score, which is shown as a black dot. The light-blue and dark-blue colors represent various sources, such as X-ray and NMR, respectively. c The ERRAT program is used to evaluate the overall model quality. It uses two horizontal lines on the error axis (representing 95% and 99% confidence levels) to show places where structural components can be reliably rejected.

**Table 2.** Assessment of the predicted three-dimensional structures of *On-*Hsp70 and *On-*Hsc70 proteins.

| Validation Index | *On-Hsp70* | *On-Hsc70* |
| --- | --- | --- |
| Ramachandran plot |  |  |
| Residues in most favored regions | 95.5% | 95.4 |
| Residues in additional allowed regions | 3.9% | 4.2% |
| Residues in generously allowed regions | 0.4% | 0.0% |
| Residues in disallowed regions | 0.2% | 0.4% |
| Overall G-factor | 0.20 | 0.20 |
| ProSA Z-Score | -11.76 | -11.96 |
| ERRAT | 92.1466 | 94.6274 |

Verify3D analysis showed an average 3D-1D score of ≥0.1 for 86.26% of *On-*Hsp70 and 80.63% of *On-*Hsc70 residues, supporting the validity of the predicted models. The ERRAT quality factor values for *On-*Hsp70 (92.1466%) and *On-*Hsc70 (94.6274%) exceeded the commonly accepted threshold of 95% for high-resolution structures.

Additional assessments included Z-scores, with values of -11.76 for *On-*Hsp70 and -11.96 for *On-*Hsc70, falling within the expected range for native proteins of similar size. The plot of residue energies revealed predominantly negative values, indicating a lack of problematic or erroneous segments in the input structures.

### Subcellular Localization

The DeepLoc-1.0 server indicated a consistent co-occurrence of both proteins and suggested a predominant presence within the cytoplasm as shown in Table 3. For *On-*Hsp70, the next most plausible subcellular locations included the nucleus, cell membrane, and extracellular space, while for *On-*Hsc70, these included the extracellular space, lysosome, and nucleus. It is noteworthy, however, that the localizations ranked as the next most probable options did not meet the server’s predefined threshold for confidence.

**Table 3.** Probability results for the subcellular localizations of a On-Hsp70 and b On-Hsc70.

| Localization | Cytoplasm | Nucleus | Extracellular | Cell membrane | Mitochondrion | Plastid | Endoplasmic reticulum | Lysosome/Vacuole | Golgi apparatus | Peroxisome |
| --- | --- | --- | --- | --- | --- | --- | --- | --- | --- | --- |
| Probability | 0.7637 | 0.3319 | 0.3223 | 0.3284 | 0.2132 | 0.0305 | 0.1050 | 0.2913 | 0.2607 | 0.1437 |

| Localization | Cytoplasm | Nucleus | Extracellular | Cell membrane | Mitochondrion | Plastid | Endoplasmic reticulum | Lysosome/Vacuole | Golgi apparatus | Peroxisome |
| --- | --- | --- | --- | --- | --- | --- | --- | --- | --- | --- |
| Probability | 0.7788 | 0.3006 | 0.3759 | 0.2958 | 0.1947 | 0.0247 | 0.1397 | 0.3111 | 0.2534 | 0.0871 |

### Protein-Protein Interaction Analysis

Table 4 revealed STRING query results indicating that stress-induced phosphoprotein 1 (stip1) stands out as the functional partner with the highest confidence score for both *On-* Hsp70 and *On-*Hsc70 proteins, with scores of 0.833 and 0.909, respectively. Among the common functional partners of *On-*Hsp70 and *On-*Hsc70, which include GAK, DNAJA4, DNAJC2, LOC100708200, LOC100703595, and Dnaja2, each plays a variety of roles in cellular processes. Interestingly, *On-*Hsp70’s unique functional partner LOC100711553 is DnaJ heat shock protein family (Hsp40) member B1a which binds to unfolded proteins while *On-*Hsc70’s bag3 is BCL2 associated athanogene 3 which not well studied yet in fish but in tetrapods and humans which has a role in chaperone binding.

**Table 4.** Predicted functional partners of *On-*Hsp70 and *On-*Hsc70.

|  | Functional Partner | Description | Score |
| --- | --- | --- | --- |
| On-Hsp70 | stip1 | Stress-induced phosphoprotein 1 | 0.883 |
|  | Hsp90ab1 | Heat shock protein 90, alpha (cytosolic), class B member 1 | 0.873 |
|  | gak | Cyclin G associated kinase | 0.863 |
|  | DNAJA4 | DnaJ heat shock protein family (Hsp40) member A4 | 0.840 |
|  | dnajc2 | DnaJ (Hsp40) homolog, subfamily C, member 2 | 0.840 |
|  | LOC100708200 | DnaJ heat shock protein family (Hsp40) member A1 | 0.840 |
|  | LOC100703595 | DnaJ heat shock protein family (Hsp40) member A2b | 0.838 |
|  | LOC100701982 | Heat shock protein 90, alpha (cytosolic), class A member 1, tandem duplicate 1 | 0.837 |
|  | Dnaja2 | DnaJ heat shock protein family (Hsp40) member A2 | 0.833 |
|  | LOC100711553 | DnaJ heat shock protein family (Hsp40) member B1a | 0.816 |
| On-Hsc70 | stip1 | Stress-induced phosphoprotein 1 | 0.909 |
|  | LOC100701982 | Heat shock protein 90, alpha (cytosolic), class A member 1, tandem duplicate 1 | 0.883 |
|  | bag3 | BCL2 associated athanogene 3 | 0.876 |
|  | gak | Cyclin G associated kinase | 0.863 |
|  | Hsp90ab1 | Heat shock protein 90, alpha (cytosolic), class B member 1 | 0.858 |
|  | dnajc2 | DnaJ (Hsp40) homolog, subfamily C, member 2 | 0.840 |
|  | DNAJA4 | DnaJ heat shock protein family (Hsp40) member A4 | 0.839 |
|  | LOC100708200 | DnaJ heat shock protein family (Hsp40) member A1 | 0.839 |
|  | LOC100703595 | DnaJ heat shock protein family (Hsp40) member A2b | 0.837 |
|  | Dnaja2 | DnaJ heat shock protein family (Hsp40) member A2 | 0.832 |

### Molecular Docking

Molecular docking was conducted for both proteins in association with ATP as their ligand demonstrated in Fig. 5. Remarkably, among the ten docking models generated for Hsp70, eight exhibited highly probable binding, evidenced by confidence scores exceeding 0.70, while the remaining two models demonstrated potential binding. On the other hand, for Hsc70, seven out of the ten models showcased robust binding, while three models displayed possible binding. This outcome accentuates the comparable binding affinities observed between the two molecular chaperones and ATP, shedding light on their shared molecular recognition mechanisms.

**Figure 5.**
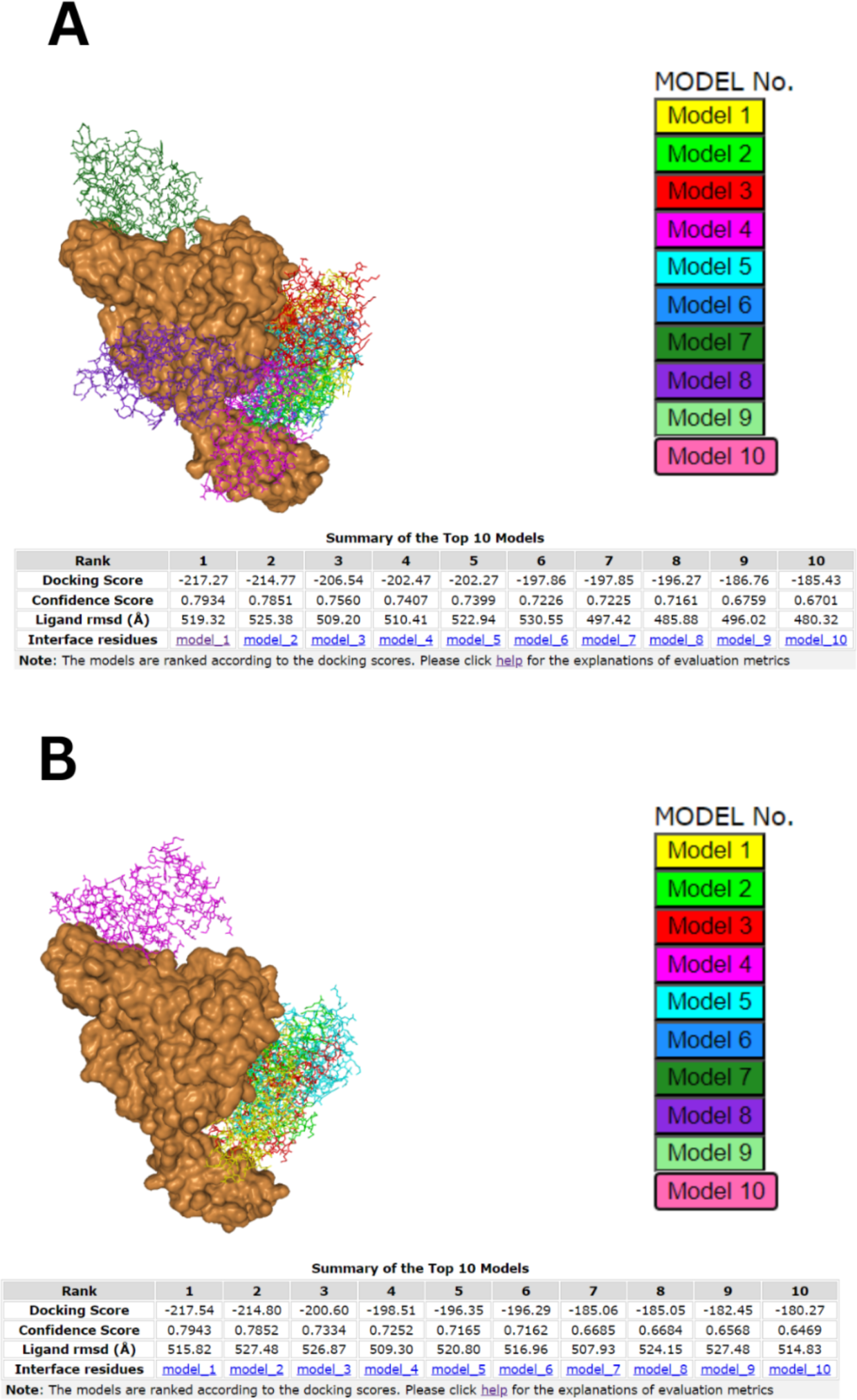
Top ten models showing binding interactions of (a) *On*-Hsp70 and (b) *On*-Hsc70 with ATP as the ligand.

### Structure similarity analysis

TM-scores >0.5 in Table 5 show that the folding of the two proteins is comparable. Notably, *On-*Hsp70, *On-*Hsc70, and the *Bos taurus* Hsc70 protein (PDB ID: 1yuwA) have a TM-score of 0.99.

**Table 5.** Top five identified structural analogs in the Protein Data Bank (PDB) library.

|  | Protein | Species | PDB Hit | TM-score | RMSDa | IDENa | Cov. |
| --- | --- | --- | --- | --- | --- | --- | --- |
| On-Hsp70 | Hsc | Bos taurus | 1yuwA | 0.99 | 1.24 | 0.892 | 1.000 |
|  | Hsp | Homo sapiens,<br>Rattus norvegicus | 4j8fA | 0.69 | 1.95 | 0.672 | 0.715 |
|  | Hsp | Homo sapiens | 3iucC | 0.68 | 1.09 | 0.684 | 0.688 |
|  | RAC | Saccharomyces cerevisiae | 7x3kB | 0.67 | 3.42 | 0.271 | 0.752 |
|  | BiP | Cricetulus griseus | 7a4uA | 0.67 | 2.27 | 0.507 | 0.701 |
| On-Hsc70 | Hsc | Bos taurus | 1yuwA | 0.99 | 1.22 | 0.923 | 1.000 |
|  | Hsp | Homo sapiens,<br>Rattus norvegicus | 4j8fA | 0.69 | 1.94 | 0.656 | 0.715 |
|  | Hsp | Homo sapiens | 3iucC | 0.68 | 1.09 | 0.695 | 0.688 |
|  | RAC | Saccharomyces cerevisiae | 7x3kB | 0.67 | 3.42 | 0.264 | 0.752 |
|  | BiP | Cricetulus griseus | 7a4uA | 0.67 | 2.27 | 0.513 | 0.701 |

### Function prediction

Results from the COFACTOR server showed that an Adenylyl Imidodiphosphate (AMP-PNP) binding site was predicted for *On-*Hsp70 based on human Hsp70 (PDB ID: 2e8aA, C-score = 0.83). In accordance with previous findings, *On-*Hsc70 has an ATP binding site as per structural similarity with bovine Hsc70 (PDB ID: 1kaxA, C-score = 0.85). Furthermore, COFACTOR predicted that the *On-*Hsp70 and *On-*Hsc70 proteins have phosphate ion (PO4)-binding sites, based on their structural similarity to other proteins with known PO4- binding sites, such as human Hsp70 proteins 3jxuA (C-score=0.42) and3gdqA (C-score=0.42), respectively.

Furthermore, in Table 6 I-TASSER’s Enzyme Commission analysis predicted that both *On-* Hsp70 and *On-*Hsc70 are structurally similar to the isozymes rhamnulokinase, glucokinase, hexokinase, and xylulokinase, all of which can transfer an inorganic phosphate group from ATP to a substrate. Predicted Gene Ontology (GO) Terms also showed that heterocyclic compound binding such as ATP binding is a molecular function of both proteins.

**Table 6.** Enzyme Commission (EC) predictions for *On-*Hsp70 and *On-*Hsc70 proteins.

|  | Enzyme | C-score | PDBHit | TM-score | RMSD | IDEN | Cov. | EC Number | Predicted Active Site Residue |
| --- | --- | --- | --- | --- | --- | --- | --- | --- | --- |
| On-Hsp70 | rhamnulokinase | 0.060 | 2uytA | 0.398 | 5.94 | 0.081 | 0.544 | 2.7.1.5 | 203,340,369 |
|  | Hexokinase glucokinase | 0.060 | 1v4tA | 0.325 | 4.87 | 0.060 | 0.411 | 2.7.1.2 2.7.1.1 | NA |
|  | xylulokinase | 0.060 | 2nlxA | 0.393 | 4.46 | 0.119 | 0.482 | 2.7.1.17 | 14,18 |
|  | glucokinase | 0.060 | 1q18A | 0.354 | 4.90 | 0.133 | 0.454 | 2.7.1.2 | NA |
|  | hexokinase | 0.060 | 3b8aX | 0.441 | 4.88 | 0.111 | 0.558 | 2.7.1.1 | NA |
| On-Hsc70 | rhamnulokinase | 0.060 | 2uytA | 0.399 | 5.96 | 0.092 | 0.544 | 2.7.1.5 | 201,338,367 |
|  | Hexokinase glucokinase | 0.060 | 1v4tA | 0.325 | 4.87 | 0.053 | 0.411 | 2.7.1.2 2.7.1.1 | NA |
|  | xylulokinase | 0.060 | 2nlxA | 0.393 | 4.46 | 0.110 | 0.480 | 2.7.1.17 | 12 |
|  | glucokinase | 0.060 | 1q18A | 0.354 | 4.91 | 0.125 | 0.454 | 2.7.1.2 | NA |
|  | hexokinase | 0.060 | 3b8aX | 0.441 | 4.89 | 0.111 | 0.558 | 2.7.1.1 | NA |

*Note.* This is a note about the table

## Discussion

### Sequence retrieval

Sequence alignment provides strong support for the claim made in the literature that Hsp70 and Hsc70 have an 85% sequence identity and are thought to play similar roles in cells (Kabani and Martineau, 2008). The present study’s high degree of sequence similarity underlines the evolutionary conservation and functional similarity of these two proteins.

This similarity in sequence identity provides more evidence that Hsp70 and Hsc70 have related and overlapping functions in cellular processes.

### Physicochemical Characteristics

Among the novel findings, the computed pI value revealed the acidic nature of the proteins, signifying their suitability for purification using isoelectric focusing on a polyacrylamide gel (pI < 7). Furthermore, the instability indicated the stability of both proteins (II < 40).

Similarly, the aliphatic index values underscored their stability across a broad temperature range. Additionally, the GRAVY confirmed that these proteins are more likely to be globular and hydrophilic rather than membranous and hydrophobic.

### Domain Architecture Analysis

The MreB and Mbl proteins from bacteria, as well as two similar archaeal sequences, make up this family. MreB is a protein that contributes to the construction of the bacterial cytoskeleton and is known to determine the rod shape in bacteria. MreB/Mbl-coding genes are exclusive to elongated bacteria and not coccoid forms (Salas et al., 2011). According to a theory, the eukaryotic cytoskeleton’s components, tubulin and actin, may have originated from prokaryotic precursor proteins linked to the modern bacterial proteins FtsZ and MreB/Mbl (Mayer, 2003). The chaperone Hsp70, the actin subfamily, and sugar kinases are all members of the broader superfamily that contains actin.

### Comparative Homology Modeling

The extensive tools for validation confirm the high quality and dependability of the suggested 3D models for *On-*Hsp70 and *On-*Hsc70. Both models satisfy strict structural requirements, as shown by the positive results of the Ramachandran plot analysis, G-factor calculation, Verify3D analysis, ERRAT quality factor evaluation, Z-scores, and residue energies plot.

These results demonstrate the homology models’ potential as accurate representations of the corresponding heat shock proteins and provide confidence in their applicability for additional research. The models’ agreement with multiple validation metrics and their alignment with experimental data support the validity of the structural predictions and open the door to further in-depth studies of the functional properties and interactions of *On-* Hsp70 and *On-*Hsc70.

### Subcellular Localization

The localization patterns observed for *On-*Hsp70 and *On-*Hsc70 are consistent with existing literature. A study by Rai et al. (2021) supports this notion as it was also found that human Hsp70 and Hsc70 were seen primarily in the cytoplasm supporting multiple metabolic processes.

Further supporting evidence comes from Balogi et al. (2019), who noted that Hsp70 is typically found in the cytoplasm but can translocate to the nucleus under stress conditions. The protein’s presence in the endosomal-lysosomal system, on the cell membrane, and in the extracellular environment during pathophysiological states adds complexity to its subcellular distribution.

For *On-*Hsc70, the findings align with studies such as Bonam et al. (2019), confirming its presence in the nucleus and external exosomes. Additionally, its localization in lysosomes is supported by its involvement in chaperone-mediated autophagy, as described by Tekirdag and Cuervo (2018). These diverse localizations underscore the multifunctional roles of Hsc70 in cellular processes beyond the cytoplasm, emphasizing its involvement in various cellular compartments and pathways. The integration of computational predictions with experimental evidence enhances our understanding of the subcellular distribution of *On-* Hsp70 and *On-*Hsc70, providing valuable insights for further functional investigations.

### Protein-Protein Interaction Analysis

The STRING analysis provided valuable insights into the potential functional partners of *On-* Hsp70 and *On-*Hsc70. Stip1, also known as stress-induced Phosphoprotein 1, is a well-known co-chaperone of heat shock proteins. This protein plays a pivotal role in facilitating protein folding and stability, particularly in times of cellular stress. Beyond its chaperone function, stip1 has been implicated in promoting cell proliferation and is even suggested to act as an oncogenic factor (Tsai et al., 2018).

Among the common functional partners of *On-*Hsp70 and *On-*Hsc70, which include GAK, Dnaja4, Dnajc2, Loc100708200, Loc100703595, and Dnaja2, each plays a variety of roles in cellular processes. GAK is associated with cyclin G and is involved in cell cycle regulation and proliferation signaling. The Dnaja4 is a member of the DnaJ heat shock protein family and contributes to cellular stress responses and protein homeostasis. The Dnaja2, another member of the DnaJ family, assists heat shock proteins in protein folding. The Loc100708200 and Loc100703595, both members of the Hsp40 class, serve as co-chaperones to ensure proper protein folding. Dnaja2, also an Hsp40 member, plays a vital role in chaperone machinery by assisting in protein folding and stability. Collectively, these proteins contribute to the cellular stress response, maintenance of protein homeostasis, and the proper functioning of proteins in the cell, especially in challenging conditions.

For unique partners, Loc100711553, in humans, are mostly found in cytoplasm and nucleus that interacts with Hsp 70 and can stimulate its ATPase activity and association between Hsc 70 and HIP while bag3 (BAG family molecular chaperone regulator 3) are found in cytoplasm, membrane, neuron projection, nucleus and plasma membrane which function as co-chaperone for Hsp 70 and Hsc 70 chaperone proteins. Acts as a nucleotide-exchange factor (NEF) promoting the release of ADP from the Hsp 70 and Hsc 70 proteins thereby triggering client/substrate protein release.

### Molecular Docking

The molecular docking results show that *On-*Hsp70 and *On-*Hsc70 have remarkably similar ATP binding affinities. Most docking models have high confidence scores, indicating strong and probable binding interactions between ATP and chaperones. The ATP-binding sites of *On-*Hsp70 and *On-*Hsc70 appear to have some functional similarities based on their shared molecular recognition mechanism.

Intriguing questions concerning the structural and functional similarities between these two heat shock proteins are also brought up by their similar binding affinities. Comprehending the subtleties of their associations with ATP can aid in clarifying their functions in cellular procedures, like the folding and unfolding of proteins dependent on ATP. These docking results lay the groundwork for additional research into the residues and molecular characteristics that influence the binding affinity of *On-*Hsp70 and *On-*Hsc70 with ATP, offering valuable insights into their functional mechanisms and potential regulatory pathways.

### Structure Similarity Analysis

The two proteins, *On-*Hsp70 and *On-*Hsc70, fold similarly when their TM-scores are more than 0.5. According to Sharma and Masison’s work (2009), the study’s results, especially the TM-scores higher than 0.66 (Table 4), demonstrate how highly conserved both proteins are across species, including humans, yeast, and rodents.

One interesting finding is that the Bos taurus Hsc70 protein (PDB ID: 1yuwA), *On-*Hsp70, and *On-*Hsc70 all have TM-scores of 0.99, which indicates almost exact structural similarities. The same Cov. Scores bolster their homology even more. The three-dimensional structures of these proteins appear to be more similar than their linear amino acid sequences, based on minor variations in the RMSD and IDEN scores. The complex folding patterns that support the overall structural conservation seen may be the source of this subtle variation.

The challenges caused by the lack of fish Hsp/Hsc70 templates in the Protein Data Bank (PDB) are also acknowledged in the discussion. The inability to compare the primary and secondary structures of *On-*Hsp70 and *On-*Hsc70 with other proteins may have resulted from this absence. These restrictions on the availability of templates highlight the need for care when interpreting some structural metrics and highlight the significance of further structural studies using templates specific to fish in order to improve our comprehension of the structural properties of these heat shock proteins.

### Function Prediction

Results from the COFACTOR server provided insights into potential ligand binding sites for *On-*Hsp70 and *On-*Hsc70. The prediction of an AMP-PNP binding site for *On-*Hsp70 suggests the presence of an ATP binding site. AMP-PNP, an ATP analog, is commonly used to study ATPases and their roles in cellular processes, further supporting the likelihood of an ATP binding site in *On-*Hsp70. Similarly, structural similarity with bovine Hsc70 confirms the presence of an ATP binding site in *On-*Hsc70. Additionally, COFACTOR predicted phosphate ion (PO4)-binding sites for both proteins, reinforcing their structural similarity to proteins with known PO4-binding sites.

Further analysis by I-TASSER revealed structural similarities between *On-*Hsp70 and *On-* Hsc70 and isozymes such as rhamnulokinase, glucokinase, hexokinase, and xylulokinase. These enzymes are known for transferring inorganic phosphate groups from ATP to substrates. Predicted Gene Ontology (GO) Terms indicated the molecular function of heterocyclic compound binding, including ATP binding, for both proteins.

These findings align with previously described operational mechanisms, where Hsp70s in their ATP-bound state rapidly capture and release their substrates, whereas Hsp70s in their ADP-bound state firmly grasp them. Hsp70s carry out their chaperone role by transitioning between the ATP and ADP-bound states. Nevertheless, the analysis lacked the sensitivity to differentiate between the roles of the constitutive (Hsc) and inducible (Hsp) isoforms. This limitation is not a significant hindrance since it has been suggested that the functional disparity primarily pertains to regulating interactions between the substrate-binding domain (SBD) and substrates rather than the inherent properties of the two ATPase domains.

With all these analyses, we have observed that similar to humans, *On-*HSP70 and *On-*HSC70 share high degree of similarities. Further, Hsp70 protein is the umbrella of molecular chaperones and folding catalysts that aid a wide range of protein folding activities within the cell which is coupled to the action of other chaperones like the Hsc70 (Mayer and Bukau, 2005). Therefore, Hsp70 proteins with Hsc70 and other cooperating chaperones constitute a complex network of folding machines that are needed to maintain cellular protein homeostasis.

## Conclusions

To gain a deeper understanding of the structural and functional biology of Hsp70 and Hsc70 in Nile tilapia, various computational tools were employed. These tools were used to analyze the physical and chemical properties, create accurate models of both proteins, and predict their functions by comparing their structural features to those of other known proteins.

Notably, the study highlighted that the generated 3D models from the study are highly reliable. Significant structural resemblance to Hsp70/Hsc70 proteins from various animal species that are taxonomically distant also supports the idea of a remarkably high degree of evolutionary conservation within this protein family. Also, the unique presence of BAG3 as functional partner of *On-*Hsc70 protein revealed additional potential diversity in the regulation of Hsc70/Hsp70 chaperones. Functional annotation based on structural similarity provides reliable indirect evidence of substantial functional conservation of these two proteins in tilapia fish. However, it is worth noting that this method may not be sensitive enough to differentiate between the two isoforms. Furthermore, the findings indicate that *On-*Hsp70 and *On-*Hsc70 have similar protein interactions and subcellular localization meaning that these two proteins are more likely to be found together, working together that makes up an organized network of folding machines.

In summary, even though the assignment of gene functions based on protein structure similarity is currently limited by the availability of protein structures in the PDB, it shows promise as a rapid, cost-effective, and reasonably reliable approach for assigning gene functions in fish on a large scale.

## Acknowledgements

The authors would like to acknowledge the extended help and expertise of Mr. Jazon Bitacura.

## Notes

### Competing Interest Statement

The authors have declared no competing interest.

